# Hybrid Neural-Cognitive Models Reveal Flexible Context-Dependent Information Processing in Reversal Learning

**DOI:** 10.1101/2025.07.31.667923

**Authors:** Yifei Cao, Maria K. Eckstein

## Abstract

Reversal learning tasks provide a key paradigm for studying behavioral flexibility, requiring individuals to update choices in response to shifting reward contingencies. While reinforcement learning (RL) models have been widely used to provide interpretable explanations of human behavior on similar learning tasks, recent work has revealed that they often fail to fully account for the complexity of learning dynamics. In contrast, artificial neural networks (ANNs) often achieve substantially higher predictive accuracy, but lack the interpretability afforded by classical RL models.

To bridge this gap, we introduce HybridRNNs—neural-cognitive models that integrate RL-inspired structures with flexible recurrent architectures. Among them, Context-ANN incorporates latent reward history and choice perseverance, and demonstrates improved alignment with human behavior compared to traditional RL models across two datasets. While not perfectly replicating human strategies, Context-ANN achieves comparable predictive accuracy to generic RNNs and offers interpretable value representations. Additional analyses of hidden dynamics reveal structured, context-sensitive internal states that adapt following reversals.

These results suggest that humans may rely on more flexible or context-sensitive learning strategies even in simple reversal tasks, and highlight the potential of HybridRNNs as cognitive models that balance interpretability and predictive accuracy.

## Introduction

The ability to learn flexibly is of vital importance in a rich and dynamic world (MacDowell et al., 2022). In cognitive neuroscience research, reversal learning tasks have been extensively used to study behavioral flexibility (Groman et al., 2019; Rudebeck et al., 2013). Unlike classical reward learning tasks, where reward contingencies remain stable, reversal learning tasks require participants to first learn the association between choices and reward outcomes over a series of trials, only for these contingencies to later be reversed. The ability to quickly adapt behavior and switch to the newly rewarding option after a reversal reflects an individual’s behavioral flexibility in reward learning (Bartolo & Averbeck, 2020).

To understand the computations underlying flexible reversal learning, researchers have developed various computational models, including reinforcement learning (RL) (Daw et al., 2011; Farashahi et al., 2017; Wilson & Collins, 2019; Eckstein et al., 2022) and Bayesian inference (Costa et al., 2015; Eckstein et al., 2022). Here, we use the term “RL” following conventions in cognitive modeling, referring to trial-by-trial learning driven by reward prediction errors. This includes classical models such as Rescorla–Wagner (Rescorla, 1972), widely adopted as a foundational RL algorithm in this context. These models update value estimates using simple rules and few parameters, enabling interpretable accounts of learning. Extensions have further improved their flexibility in reversal tasks by incorporating adaptive learning rates or value resets (Hauser et al., 2014; Sidarus et al., 2019; Barnby et al., 2022). Howeverever, simple, hand-crafted models often fail to fully capture behavioral data (Peterson et al., 2021; Miller et al., 2024; Eckstein et al., 2024). Moreover, their reliance on predefined assumptions can introduce biases and researcher subjectivity, potentially leading to inaccurate or incomplete representations of behavior (Feher da Silva & Hare, 2020).

An alternative approach to understanding cognitive computations is to discover the underlying cognitive strategies automatically from observed behavior. One such approach involves modeling human behavior using artificial neural networks (ANNs). Compared to classical cognitive models, ANNs require no handcrafted assumptions and structural hypotheses while offering the flexibility to approximate a wide range of models by adjusting their weights. Previous studies have successfully applied ANNs to model human behavior in decisionmaking (Dezfouli et al., 2019), reward learning (Song et al., 2021), and cognitive flexibility (Jaffe et al., 2023), amongst others, achieving higher predictive accuracy in forecasting future behavior. However, the large number of parameters in ANNs poses a major interpretability challenge, necessitating additional efforts to uncover the cognitive and computational strategies they employ.

To achieve both predictive accuracy and interpretability in modeling learning process, recent techniques have been developed such as DisRNN (Miller et al., 2024), Tiny RNN (Ji-An et al., 2023), and HybridRNN (Eckstein et al., 2024). We use the HybridRNNs approach here, which integrates the flexibility of ANNs with the interpretable structure of RL models. Specifically, HybridRNN (Eckstein et al., 2024) replaces the hand-crafted Q-value updating function and choice perseverance mechanisms with neural networks, hereby allowing the model to discover the arbitrary functions that underlie these processes. In this way, HybridRNN allows the incorporation of additional cognitive factors relevant to reward learning, such as context information (Palminteri et al., 2015) and various types of memory that go beyond RL (Collins & Frank, 2012; Davidow et al., 2016; Gershman & Daw, 2017), as inputs to assess their impact on model performance. This framework enables an automated identification of the cognitive mechanisms that support human flexible learning under different conditions.

In this study, we fitted HybridRNN models to human reversal learning data. We first compared classical RL models and artificial neural networks (ANNs), identifying a clear gap in predictive accuracy. To address this, we developed a series of HybridRNNs by replacing hand-crafted value update rules with learnable neural components. Among these, the Context-ANN—incorporating choice perseverance and context inputs—achieved the highest predictive accuracy.

We further conducted qualitative comparisons and found that Context-ANN more closely resembled human behavior during reversals than other models. Analysis of its value updating channel revealed a context-dependent updating strategy. Additionally, we examined the model’s hidden dynamics, identifying structured population activity that adapts to task changes, including distinct attractor-like clusters and reversal-aligned transitions. Together, these findings highlight the utility of HybridRNNs for both fitting behavior and probing the internal computations that support flexible learning.

## Methods

### Experimental Design and Dataset

The reversal learning task is a paradigm designed to assess an individual’s ability to adapt their behavior in response to a dynamically changing reward environment. In our task, participants were presented with two choices, *C*_1_ and *C*_2_, each associated with continuous reward *r*_1_ and *r*_2_. One choice initially offers a larger reward magnitude than the other for several trials. However, in the current task, at unpredictable intervals and without explicit cues, the reward contingencies reverse. Following this reversal, the previously more rewarding choice becomes the less rewarding one, requiring participants to adjust their decision-making strategies accordingly.

The present study analyzed human behavioral performance using an open-access dataset from Lee et al. (2023). In their two-armed bandit task, the mean reward distributions associated with each option followed a random walk over time. The mean rewards, 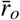, for the two options *o* ∈ {1, 2} were symmetrical, such that 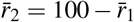. Reversals were defined as the time points where the mean rewards for the two options intersected (see Fig. 1C). Following the filtering criteria outlined in Lee et al. (2023), we included 669 sessions of data, each consisting of 80 trials. To further examine the generalizability of our findings, we also analyzed another independent reversal learning dataset from Eckstein et al. (2022). Detailed results and methods are provided in the Supplementary Materials.

**Figure 1:**
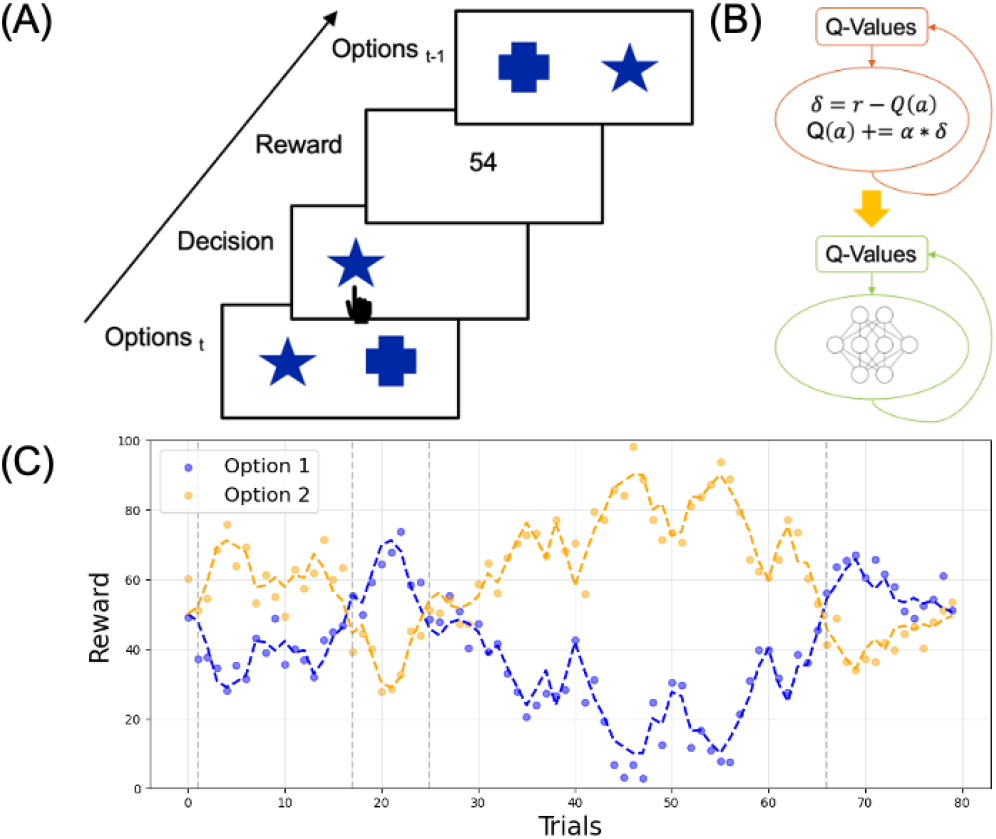
Task design and value updating in HybridRNNs. (A) On each trial, participants choose between two options and receive feedback. (B) HybridRNNs replace hand-crafted RL update rules with a neural network to learn value updates. (C) Reward schedule over 80 trials. Colored dots show actual rewards for each option, and dashed lines indicate the underlying mean rewards. Vertical gray lines mark reversal points.

### Model Architectures

#### Reinforcement Learning Models

To identify the best classic cognitive model, we systematically compared reinforcement learning models with different structures (Wilson & Collins, 2019). The Rescorla-Wagner (RW) model assumes that learning is driven by unexpected outcomes, including surprising occurrence or omission of reward during associative learning (Rescorla, 1972). In a RW model, the prediction error (PE) signal defines the difference between observed and expected reward. This PE signal is weighted by a learning rate parameter when individuals update their reward expectations. According to this model, the agent’s expected reward for the chosen action on trial *t, Q*(*t*) is calculated as follows:

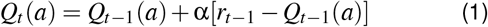

In this equation, *Q*_*t*_(*a*) refers to the reward value expectation for the chosen option at trial *t, Q*_*t*−1_ is the reward value expectation for the chosen option at trial *t*− 1, the *r*_*t*−1_ (given that rewards were normalized to 0 and 1) refers to the reward actually perceived by participants at trial *t*− 1, and the α represents the learning rate. We call this processing step the “value channel” because it processes reward-related value information.

At trial *t*, for action selection, the vector *Q*(*t*) of all action values is transformed into a vector of choice probabilities *p*_*t*_ of the same length using the softmax function. The transformation is determined by a free parameter “inverse decision temperature” β, to induce more deterministic or more random choices:

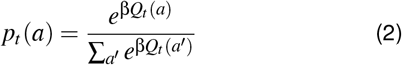

For increasingly flexible RL models, we followed the work of Eckstein et al. (2022) to include counterfactual value updating and choice persistence into the framework. With counterfactual value updating, in each trial *t*, the value of the unchosen action is also updated, using an “imaginary” counterfactual reward that is the inverse of the actually received reward 1− *r*_*t*−1_ (given that rewards were normalized to 0 and 1), and based on the same learning rate as the chosen action:

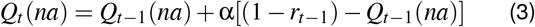

In this equation, *Q*_*t*_(*na*) refers to the reward value expectation for the unchosen option at trial *t*. We call this additional, parallel processing step the “counterfactual value channel”.

Finally, we extended the RL models with “perseverance”, which enables action repetition independently of rewards. The “perseverance” term *c* adds a small “bonus” (of size κ) to the value of the action *a* that was chosen on the previous time step, but not to all other actions *na*:

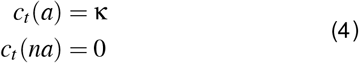

#### RL-ANN Architecture

The general idea of creating RL-ANN is to replace the value and perseverance updating function from Best RL with neural networks. For the three ‘channels’ designed in classic RL models, their hand-crafted updating functions (see Equations (1,2,3)) were replaced with a simple neural network.

For each trial, RL-ANN’s value channel takes the reward *r*_*t*−1_ and the value of chosen action *Q*_*t*−1_ from the previous trial as inputs, just like the Simple RL model:

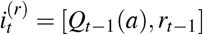

The hidden layer activations, referred to as the “state” 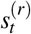, are computed by processing the input vector through the network’s first fully-connected layer. Specifically, the input vector is multiplied by the weight matrix 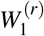, the bias term 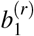 is added, and the result is passed through a tanh activation function to introduce non-linearity:

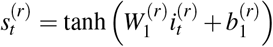

The output of value channel is the update to the value of chosen action *Q*_*t*_(*a*), and is calculated by passing the state 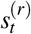 through the second fully-connected layer with weights 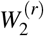 and biases 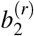

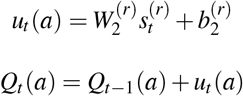

The only difference between Simple RL and RL-ANN’s value channels hence is that we replaced the linear updating function from Simple RL to neural networks in RL-ANN.

We next turned to the counterfactual value channel, which has the same structure as the value channel: it takes previous reward *r*_*t*−1_ and value of unchosen action *Q*_*t*−1_(*na*) as input and predicts the update to the current value of the unchosen action:

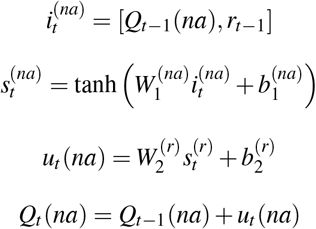

Hence, RL-ANN’s counterfactual channel has the same structure as Simple RL’s.

The perseverance channel is also a 3-layer fully-connected multilayer perceptron (MLP). The input to the channel is the action from the previous trial *a*_*t*−1_, and the output of the channel is a vector *c*_*t*_ ∈ ℝ^2^. For binary choice tasks, each dimension of *c*_*t*_ represents the perseverance scalar corresponding to one of the two available actions.

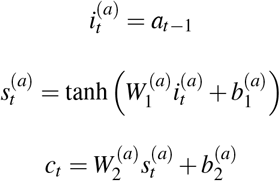

Finally, depending on the need to combine the channels, the values *Q*_*t*_(*a*), counterfactual values *Q*_*t*_(*na*), and perseverance *C*_*t*_ are combined additively and then pass through the softmax to select an action for the next trial:

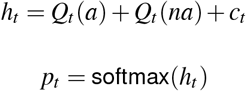

#### Context-ANN Architecture

Context-ANN extends RL-ANN by adding additional context information as input to the network, and is the winning model in the current paper. We provide context information for value channel with the vector *Q*_*t*−1_ and perseverance channel with the vector *c*_*t*−1_ as additional inputs, giving rise to access of value and perseverance of all possible choices (Palminteri et al., 2015).

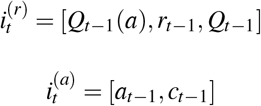

Besides the difference in model inputs, the model structure and outputs stay the same with RL-ANN.

#### Memory-ANN Architecture

In another approach of extending RL-ANN, we added states of hidden units from previous trial 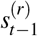 and 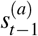 as additional inputs. This modification of input enables the model to access its own compressed representation of the past history.

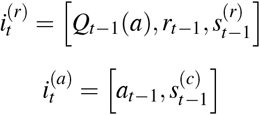

#### Vanilla RNN architecture

Vanilla RNN is a simple, fully-connected recurrent neural network with three layers. On each trial *t*, it receives the previous action *a*_*t*−1_ (one-hot encoded) and reward *r*_*t*−1_ as input. The hidden state *s*_*t*_ is updated based on the current input and the previous hidden state. This hidden state is then passed through an output layer to generate action logits, which are converted into probabilities using a softmax function:

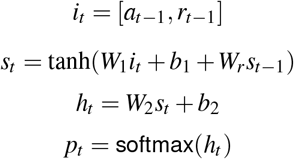

### Neural Network Training

We divided the dataset into three parts: 80% for training (537 sessions), 10% for testing (66 sessions), and 10% for validation (66 sessions). The training data was used to fit model parameters across various configurations of model weights and biases. We assessed models’ final model fitting by calculating the fit on the held-out testing data, ensuring they did not overfit to the training data.

The networks were trained in Haiku using the Adam optimizer with a learning rate of 0.005. Training was performed on batched data, incorporating a cross-entropy loss and L2-regularization on recurrent weights (with coefficients 1e-5). The maximum number of training step we used was 200,000. Early stopping was applied if, within the most recent 200 training steps, the mean validation loss of the last 50 steps did not improve compared to the first 50 steps.

### Model Fitting Objective

Models were trained to replicate human behavior (rather than to optimize task performance), by minimizing the negative log-likelihood loss of the training data under the model (cross-entropy):

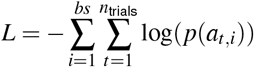

where *bs* is the batch size and *n*_trials_ = 80.

The best model for each configuration of hyperparameters was evaluated on test data, with losses converted to trial-wise prediction accuracy:

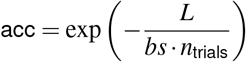

Model performance was assessed using the Akaike Information Criterion (AIC):

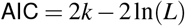

where *k* is the number of parameters. The AIC is a standard measure of model fit that balances model fit and complexity.

### Qualitative Model Comparison

Besides quantitative model fitting, we also wanted to make sure that the trained models behaved similar to human participants. To this end, we performed several behavioral analyses to test the similarity between human and model behaviors.

#### Model Staying Behavior

We analyzed the probability of repeating the previous action (*a*_*t*_ = *a*_*t*+1_), which we refer to as “staying behavior”. This probability was calculated across four reward histories, defined by combinations of current and previous rewards (*R*_*t*−1_ and *R*_*t*_), and we binned the data with reward larger or smaller than 50.

For each condition, the staying probabilities were computed for human participants and each model. These probabilities were averaged across participants, and paired *t*-tests were conducted to compare the behavior of human participants and models in each combination of previous and current rewards.

#### Model Inspection

To investigate how the reward module maps mapped rewards *r*_*t*−1_ onto values *Q*_*t*+1_(*a*), we probed the module with a range of input values and analyzed its output. Input values were uniformly sampled within defined ranges. *r*_*t*−1_ values were sampled between 1 and 100, and *Q*_*t*−1_(*a*) values were sampled between the 10th and 90th percentiles of observed values. The relevant parameters 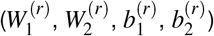 were extracted from the trained RL-ANN and Memory-ANN models. A new MLP was initialized with the same architecture as the original reward module (e.g., for Memory-ANN: 2 input units, 16 hidden units, and 1 output unit, for Context-ANN: 2 input units, 8 hidden units, and 1 output unit) for further analysis (see Table4 for details of each model).

#### Hidden Unit Dynamics Analysis

We extracted hidden state activations from the trained models by running them on the full dataset, aligning them to reversal trials. To reduce dimensionality and visualize temporal dynamics, we applied PCA to the concatenated hidden states across all trials and sessions. The first three principal components were used for trajectory visualization and condition-based comparisons (see Supplementary Methods for details).

## Results

### Quantitative Model Comparison

We first modeled the dataset using two extremes: the simplest RL model with only α and β, and a generic Vanilla RNN. Predictive accuracy improved from Simple RL (Normalized Likelihood = 63.5%, AIC = 4880.51) to Vanilla RNN (67.0%, AIC = 4533.33), indicating that classical models missed to capture predictable and systematic behavioral patterns. This highlights the flexibility of neural networks in modeling the complexity of reversal learning.

Although Vanilla RNN achieved high predictive accuracy in modeling human behavior, its large number of fitted parameters makes interpretation challenging. This raises the question: what features contribute to its higher predictability? One possibility is that the value updating rule in Simple RL is overly rigid. A traditional approach would involve manually refining the update function (Hauser et al., 2014; Barnby et al., 2022), but we instead replaced the entire update process with an ANN, implicitly testing a broad range of possible modifications.

To improve model interpretability while maintaining high predictive accuracy, we developed a series of cognitive and hybrid models by gradually adding computations (see Table 1). Compared to Simple RL, its hybrid counterpart RL-ANN (see Fig. 2A) improved predictive accuracy (*AIC*_*difference*_ = 117.01), suggesting that a more flexible value update function is necessary. Since prior research indicates that humans update the values of unchosen actions (Palminteri et al., 2016; Eckstein et al., 2022), we introduced counterfactual updating to both Simple RL and RL-ANN. Even though it had small effect in simple RL (*AIC*_*difference*_ = 37.12), it did in RL-ANN (*AIC*_*difference*_ = 223.42), showing that human learners adjust the values of choices whose outcomes they have not directly observed. Next, we incorporated choice perseverance, a widely used extension in RL models (Eckstein et al., 2022) (see Fig. 2B), and found that it enhanced prediction in both cognitive (*AIC*_*difference*_ = 248.11) and hybrid models (*AIC*_*difference*_ = 86.36). Finally, we examined whether combining counterfactual updating and choice perseverance would further improve performance. Results showed that this combination maximized the RL model’s predictability (*AIC*_*difference*_ = 350.52) but did not significantly improve RLANN beyond counterfactual updating alone (*AIC*_*difference*_ = −80.79).

**Table 1:** Quantitative model comparisons of cognitive and hybrid model series. Best-performing models are highlighted in bold.

| Classic Model | AIC Score | Normalized Likelihood | N Params. | Hybrid Model | AIC Score | Normalized Likelihood | N Params. |
| --- | --- | --- | --- | --- | --- | --- | --- |
| Simple RL | 4880.51 | 63.5% | 2 | RL-ANN | 4763.50 | 64.7% | 33 |
| RL with CU<sup>1</sup> | 4843.39 | 62.3% | 3 | **RL-ANN with CU** | **4540.08** | **66.4%** | **75** |
| RL with CP<sup>2</sup> | 4632.40 | 64.9% | 2 | RL-ANN with CP | 4677.14 | 65.6% | 66 |
| **RL with CU CP** | **4529.99** | **65.6%** | **3** | RL-ANN with CU CP | 4620.87 | 66.4% | 108 |

<sup>1</sup> CU refers to counterfactual updating. <sup>2</sup> CP refers to choice perseverance.

**Figure 2:**
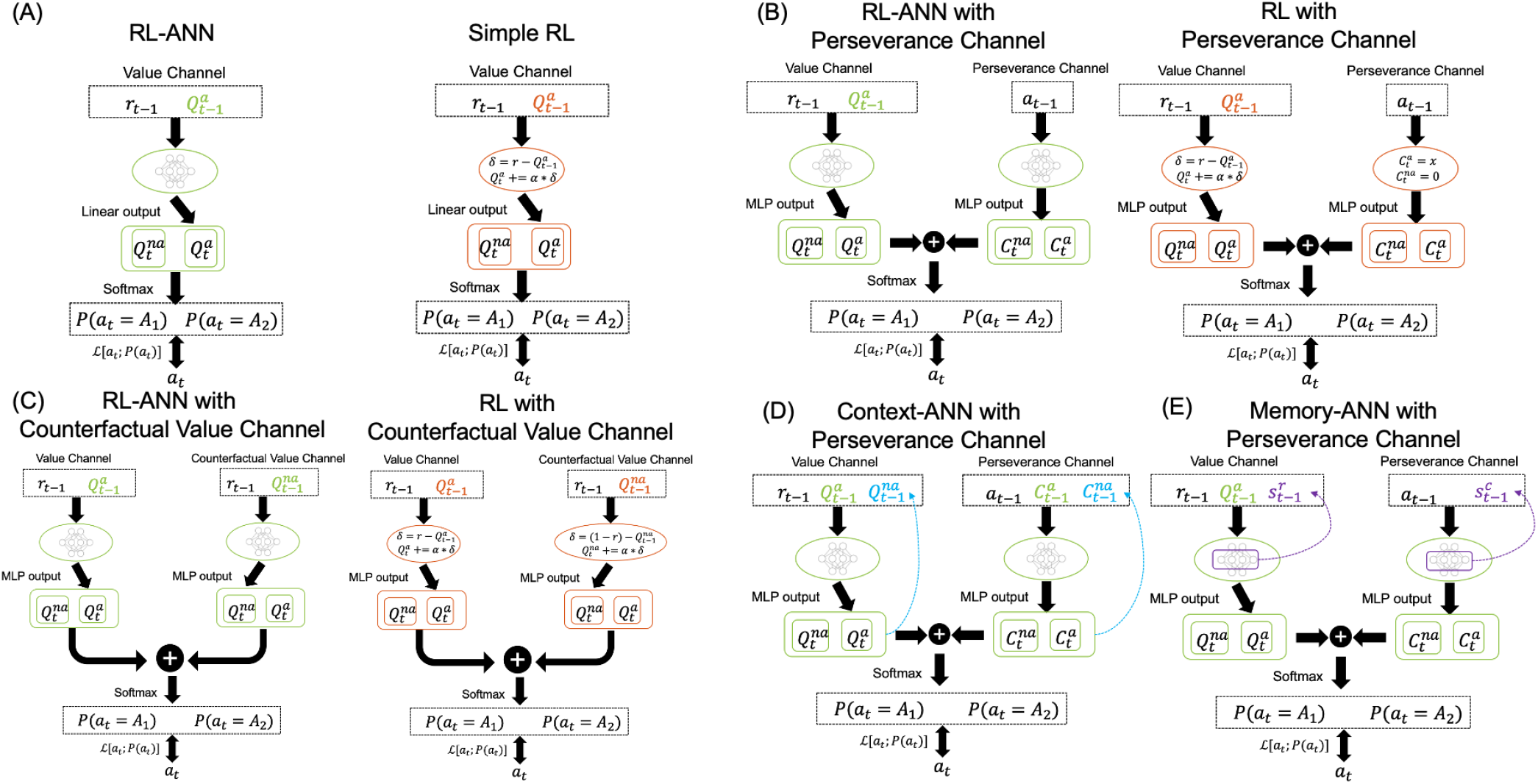
Model architectures of classic cognitive models and HybridRNN models.

Since none of these models reached the predictability of Vanilla RNN (see Table 2), we investigated whether hybrid models benefited from additional context or memory information. Model comparisons identified Context-ANN with a perseverance channel (see Fig. 2D) as the best-performing model, showing the most similar predictive accuracy with Vanilla RNN (*AIC*_*difference*_ = 54.56). Taken together, quantitative model comparisons suggest that Context-ANN achieves among the highest predictive performance across all cognitive and HybridRNN models while offering greater interpretability than Vanilla RNN; this finding is further supported by results on the independent Eckstein dataset (see Table 6 and Table 7).

**Table 2:** Quantitative model comparisons of cognitive models, RNN, and HybridRNN models. Best-performing model is highlighted in bold.

| Model name | Normalized Likelihood | AIC Score |
| --- | --- | --- |
| Simple RL | 63.5% | 4880.51 |
| Best RL<sup>1</sup> | 65.6% | 4529.99 |
| RL-ANN | 65.7% | 4664.80 |
| Memory-ANN | 66.3% | 5733.76 |
| **Context-ANN** | **67.3%** | **4478.77** |
| Vanilla RNN | 67.0% | 4533.33 |

<sup>1</sup> Best RL refers to the RL model with CU CP.

### Qualitative Model Comparison

Besides quantitative model comparisons, it is also important to qualitatively examine whether model’s behavior pattern align with human choices (Palminteri et al., 2017). We first calculated the mean reward received by each model agent and actual human behavior around reversal trials (see Fig. 3A). Although none of the models perfectly matched human performance, both Context-ANN and Vanilla RNN exhibited reward trajectories that were qualitatively most similar to human behavior at the point of reversal. Possible reasons for the remaining deviations are discussed in the Discussion section.

**Figure 3:**
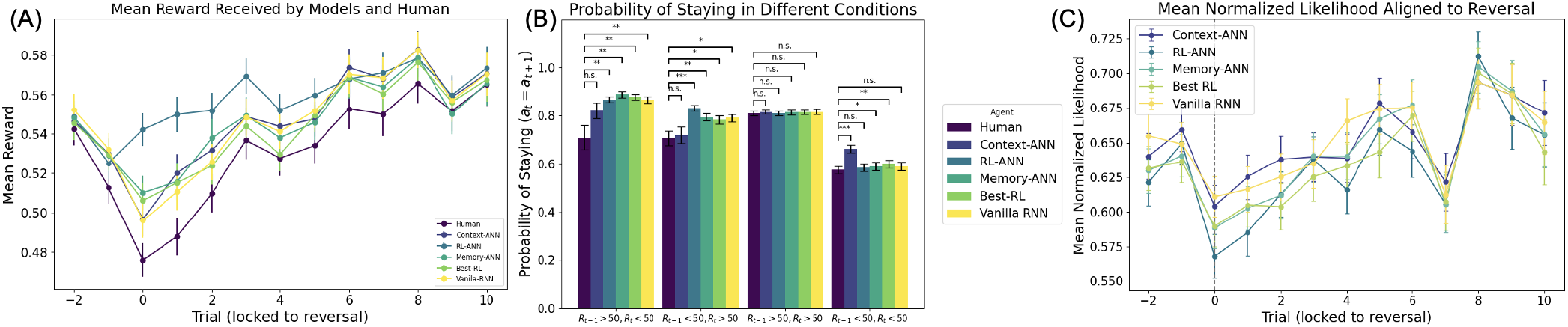
Qualitative model comparison results. (A) Mean rewards received by human and each model around the reversal trials, the reward at chance level is 0.5. (B) Probability of staying behavior of human choices and model behaviors, t-test was performed to compare between human and each model. (C) Mean normalized likelihood of each model on human held-out data, aligned to reversal trials. Context-ANN and Vanilla RNN show the largest predictive accuracy.

We then calculated the “staying behavior” (see Methods section) of each model and actual human behavior, and performed pairwise t-tests between model behavior and human choices. The result showed that, although all models showed significant difference with human choices in at least one reward condition, Context-ANN (with perseverance channel) showed relatively smaller deviations compared to other models. Notably, Context-ANN overestimated the probability of staying when no reward was received for two consecutive trials (see Fig. 3B). Taken together, although some discrepancies remain, Context-ANN emerges as the most promising model based on its overall balance between predictive accuracy and alignment with key qualitative patterns of human behavior.

We also computed trial-wise normalized log-likelihood (NLL) aligned to reversals (see Fig. 3C). Context-ANN and Vanilla RNN achieved the highest overall predictive accuracy, even though both showed a transient drop in performance following reversals.

### Information Processing in Context-ANN

To examine the internal learning dynamics of Context-ANN, we systematically simulated value updates across a range of rewards and action values (*r*_*t*_, *Q*_*t*_(*a*), *Q*_*t*_(na)) and observed the corresponding outputs *Q*_*t*+1_(*a*). While traditional RL models update action values linearly based on the reward prediction error (see Fig. 4A), Context-ANN displayed a non-linear reward-to-update mapping (see Fig. 4B), indicating a more flexible learning strategy. This non-linearity was further modulated by the value of the unchosen option: when *Q*_*t*_(*na*) was high, updates to the chosen action value were attenuated; when *Q*_*t*_(*a*) was low, the model responded more strongly to reward signals.

**Figure 4:**
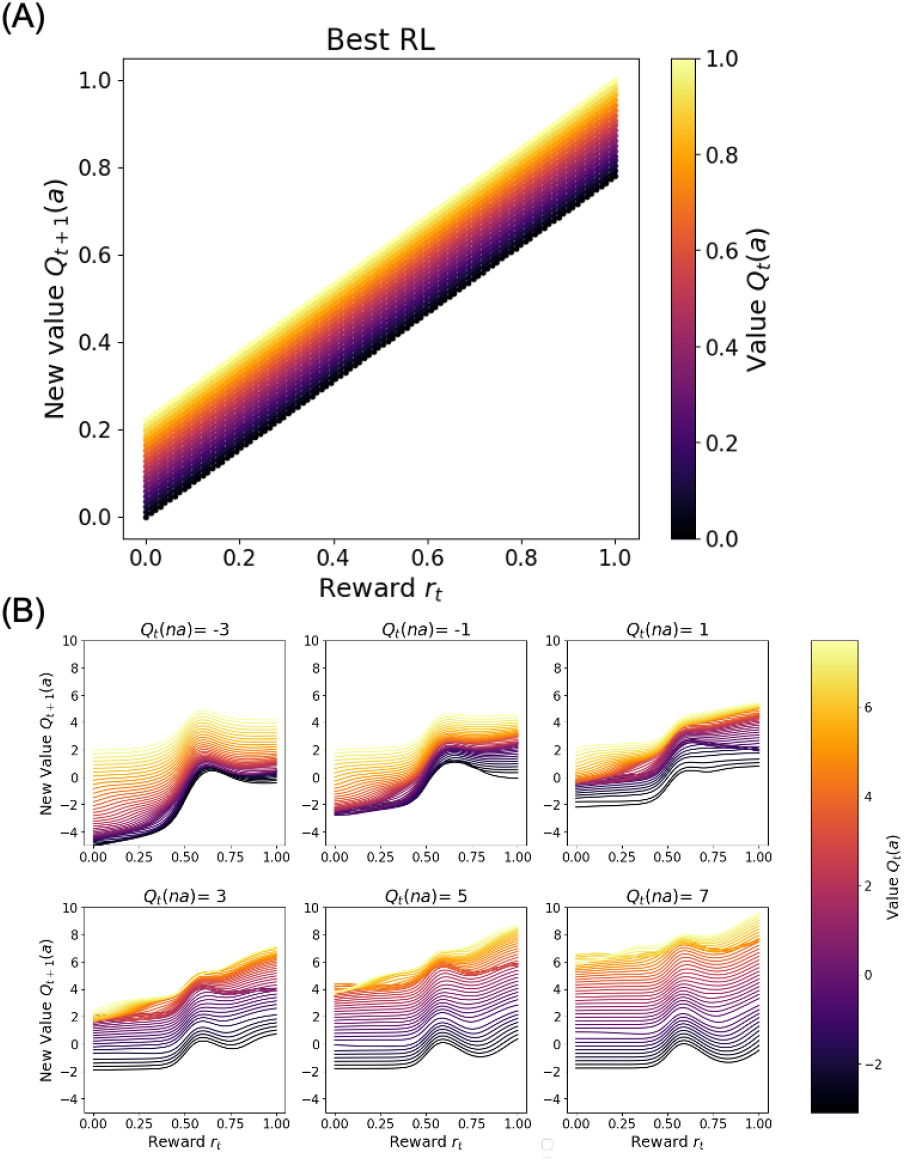
Value updating functions of Best RL and Context-ANN. (A) Linear updating function of classical RL model. (B) Nonlinear and context dependent value updating functions of Context-ANN. Note that the value scale differs between (A) and (B) because Context-ANN does not constrain value outputs within the [0, 1] range.

Additionally, we observed a sharp inflection in the update function around *r*_*t*_ = 0.5, suggesting heightened sensitivity to reward values near this threshold. This aligns with the task structure, where rewards around 0.5 indicate increased volatility and potential reversal. Such selective responsiveness would be adaptive, helping the model detect changes in the environment. Furthermore, the slope of the value update function varied systematically with *Q*_*t*_(*na*): when the unchosen action had low value, low rewards were grouped together; when *Q*_*t*_(*na*) was high, updates became more sensitive to high rewards. This pattern supports the interpretation that value updating in Context-ANN is modulated by the context defined by the alternative option’s value, consistent with prior work on context-sensitive learning strategies (Palminteri et al., 2015).

To quantify these effects, we conducted a linear regression predicting *Q*_*t*+1_(*a*) from *r*_*t*_, *Q*_*t*_(*a*), *Q*_*t*_(*na*), and their interactions. The most pronounced effect was the negative interaction between *r*_*t*_ and *Q*_*t*_(*na*) (β = − 0.32, *p <* .001), indicating that rewards were weighted more heavily when the alternative action had lower expected value. This context-dependent modulation deviates from the fixed-error structure of classical RL and supports the view that Context-ANN approximates human-like flexible learning by integrating choice context into its value updating process.

### Hidden Unit Dynamics Around Reversal

Context-ANN exhibited distinct internal dynamics aligned to reversals. Two hidden units significantly increased activation following switch points (see Fig. 5A), with ramping patterns observed around the reversal (see Fig. 5B). The analysis on the extracted PCs confirmed there is a populational response to the switch points with the first PC showing decreased score after switching (see Fig. 5C and D). Furthermore, to assess whether the hidden state encodes contextual value information, we trained a classifier to decode whether *Q*_left_ *> Q*_right_ from the hidden units. This yielded a 5-fold cross-validated accuracy of 0.788 using the hidden units and 0.764 using the top 3 principal components.

**Figure 5:**
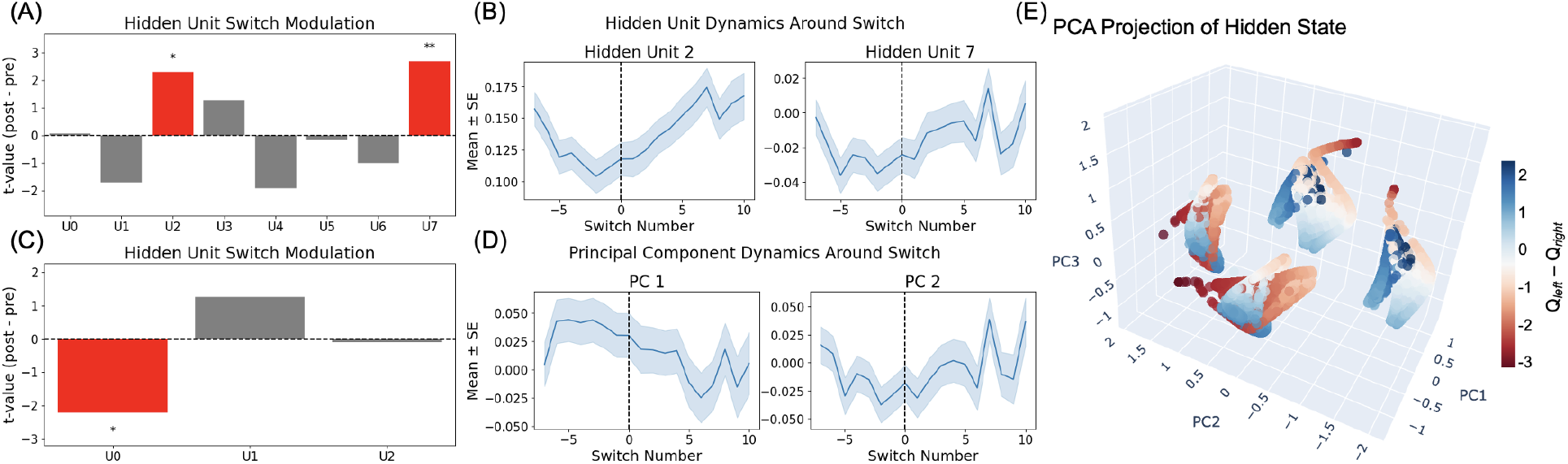
Internal dynamics of Context-ANN aligned to reversal points. (A, C) T-values for each hidden unit comparing activation before and after reversals, with significant modulations highlighted in red. (B) Temporal dynamics of example hidden units around the switch point. (D) Principal components of hidden state activity show systematic changes around reversals. (E) 3D PCA projection of hidden states reveals four attractor-like clusters, with color indicating the difference in action values (*Q*_left_™*Q*_right_).

PCA projection of the hidden states revealed four attractor-like clusters in a low-dimensional space (see Fig. 5E and Fig. 6A). Within each cluster, hidden state positions varied smoothly with the value difference between actions (*Q*_left_ − *Q*_right_), suggesting context-sensitive internal organization. Following reversals, the probability of the hidden state leaving its current attractor increased (see Fig. 7B; Table 5), indicating adaptive reconfiguration of internal dynamics in response to changing environments. These results suggest that Context-ANN maintains structured, adaptive internal representations that flexibly reorganize in response to reversals.

**Table 3:** Linear regression results on predicting value of chosen action.

| Predictor | Estimate | Std.Err | t value | p-value |
| --- | --- | --- | --- | --- |
| $r_t$ | 4.69 | 0.025 | 190.40 | <0.001 |
| $Q_t(a)$ | 0.43 | 0.003 | 122.58 | <0.001 |
| $Q_t(na)$ | 0.49 | 0.004 | 149.65 | <0.001 |
| $r_t * Q_t(a)$ | 0.04 | 0.006 | 6.20 | <0.001 |
| $r_t * Q_t(na)$ | -0.32 | 0.005 | -61.51 | <0.001 |
| $Q_t(a) * Q_t(na)$ | 0.04 | 0.001 | 84.54 | <0.001 |

**Table 4:** Model size comparison for all candidate models.

| Model | Units | N Params. |
| --- | --- | --- |
| RL-ANN | 8 | 33 |
| RL-ANN with CU | 8 | 75 |
| RL-ANN with CP | 16 | 66 |
| RL-ANN with CU CP | 8 | 108 |
| Context-ANN | 8 | 107 |
| Memory-ANN | 16 | 659 |
| Vanilla RNN | 8 | 114 |

**Table 5:**
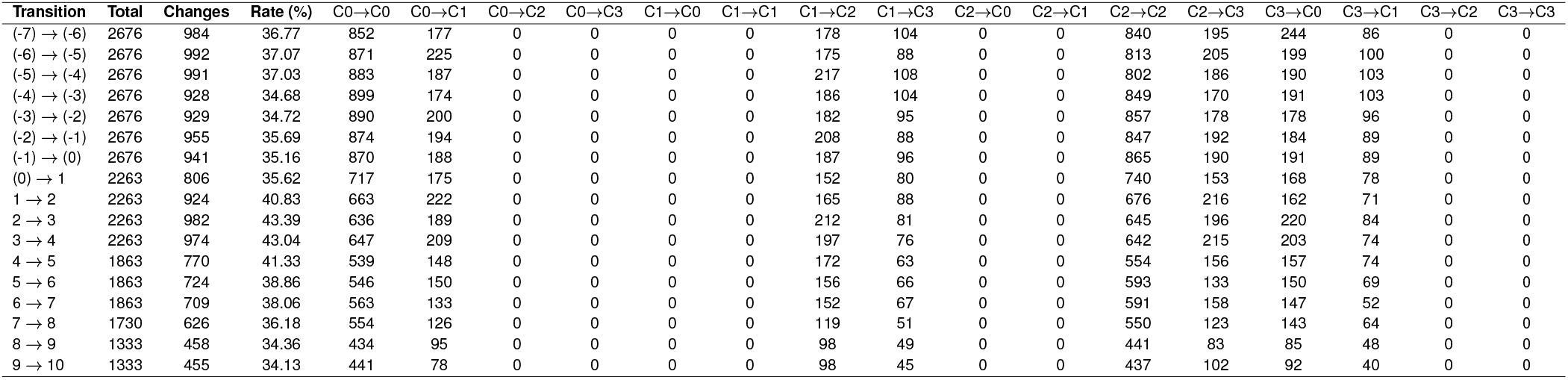
Detailed cluster transitions across different switch stages. Each cell indicates the number of transitions from one attractor to another during the specified switch transition.

| Transition | Total | Changes | Rate (%) | C0→C0 | C0→C1 | C0→C2 | C0→C3 | C1→C0 | C1→C1 | C1→C2 | C1→C3 | C2→C0 | C2→C1 | C2→C2 | C2→C3 | C3→C0 | C3→C1 | C3→C2 | C3→C3 |
| --- | --- | --- | --- | --- | --- | --- | --- | --- | --- | --- | --- | --- | --- | --- | --- | --- | --- | --- | --- |
| (-7) → (-6) | 2676 | 984 | 36.77 | 852 | 177 | 0 | 0 | 0 | 0 | 178 | 104 | 0 | 0 | 840 | 195 | 244 | 86 | 0 | 0 |
| (-6) → (-5) | 2676 | 992 | 37.07 | 871 | 225 | 0 | 0 | 0 | 0 | 175 | 88 | 0 | 0 | 813 | 205 | 199 | 100 | 0 | 0 |
| (-5) → (-4) | 2676 | 991 | 37.03 | 883 | 187 | 0 | 0 | 0 | 0 | 217 | 108 | 0 | 0 | 802 | 186 | 190 | 103 | 0 | 0 |
| (-4) → (-3) | 2676 | 928 | 34.68 | 899 | 174 | 0 | 0 | 0 | 0 | 186 | 104 | 0 | 0 | 849 | 170 | 191 | 103 | 0 | 0 |
| (-3) → (-2) | 2676 | 929 | 34.72 | 890 | 200 | 0 | 0 | 0 | 0 | 182 | 95 | 0 | 0 | 857 | 178 | 178 | 96 | 0 | 0 |
| (-2) → (-1) | 2676 | 955 | 35.69 | 874 | 194 | 0 | 0 | 0 | 0 | 208 | 88 | 0 | 0 | 847 | 192 | 184 | 89 | 0 | 0 |
| (-1) → (0) | 2676 | 941 | 35.16 | 870 | 188 | 0 | 0 | 0 | 0 | 187 | 96 | 0 | 0 | 865 | 190 | 191 | 89 | 0 | 0 |
| (0) → 1 | 2263 | 806 | 35.62 | 717 | 175 | 0 | 0 | 0 | 0 | 152 | 80 | 0 | 0 | 740 | 153 | 168 | 78 | 0 | 0 |
| 1 → 2 | 2263 | 924 | 40.83 | 663 | 222 | 0 | 0 | 0 | 0 | 165 | 88 | 0 | 0 | 676 | 216 | 162 | 71 | 0 | 0 |
| 2 → 3 | 2263 | 982 | 43.39 | 636 | 189 | 0 | 0 | 0 | 0 | 212 | 81 | 0 | 0 | 645 | 196 | 220 | 84 | 0 | 0 |
| 3 → 4 | 2263 | 974 | 43.04 | 647 | 209 | 0 | 0 | 0 | 0 | 197 | 76 | 0 | 0 | 642 | 215 | 203 | 74 | 0 | 0 |
| 4 → 5 | 1863 | 770 | 41.33 | 539 | 148 | 0 | 0 | 0 | 0 | 172 | 63 | 0 | 0 | 554 | 156 | 157 | 74 | 0 | 0 |
| 5 → 6 | 1863 | 724 | 38.86 | 546 | 150 | 0 | 0 | 0 | 0 | 156 | 66 | 0 | 0 | 593 | 133 | 150 | 69 | 0 | 0 |
| 6 → 7 | 1863 | 709 | 38.06 | 563 | 133 | 0 | 0 | 0 | 0 | 152 | 67 | 0 | 0 | 591 | 158 | 147 | 52 | 0 | 0 |
| 7 → 8 | 1730 | 626 | 36.18 | 554 | 126 | 0 | 0 | 0 | 0 | 119 | 51 | 0 | 0 | 550 | 123 | 143 | 64 | 0 | 0 |
| 8 → 9 | 1333 | 458 | 34.36 | 434 | 95 | 0 | 0 | 0 | 0 | 98 | 49 | 0 | 0 | 441 | 83 | 85 | 48 | 0 | 0 |
| 9 → 10 | 1333 | 455 | 34.13 | 441 | 78 | 0 | 0 | 0 | 0 | 98 | 45 | 0 | 0 | 437 | 102 | 92 | 40 | 0 | 0 |

**Figure 6:**
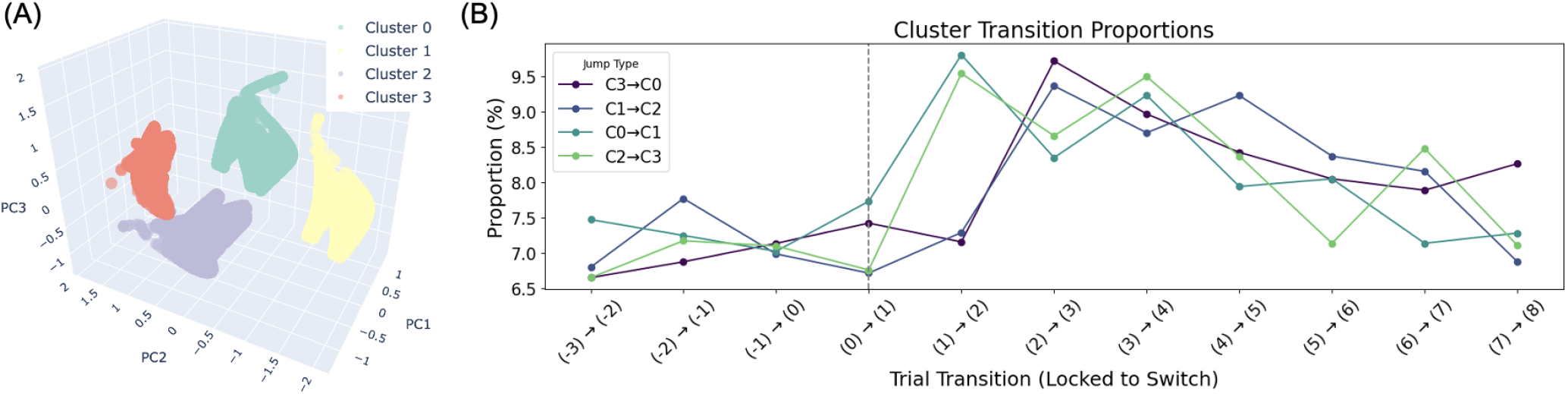
(A) PCA projection of hidden states reveals four distinct attractor-like clusters. Each cluster reflects a recurring internal representation pattern used by the network. (B) Proportions of cluster-to-cluster transitions aligned to reversal trials. Certain transition types (e.g., C0→C1, C2→C3) increase immediately after the reversal, while others (e.g., C1→C2, C3→C0) rise with a slight delay, indicating temporally specific reorganization of internal dynamics in response to environmental changes.

**Figure 7:**
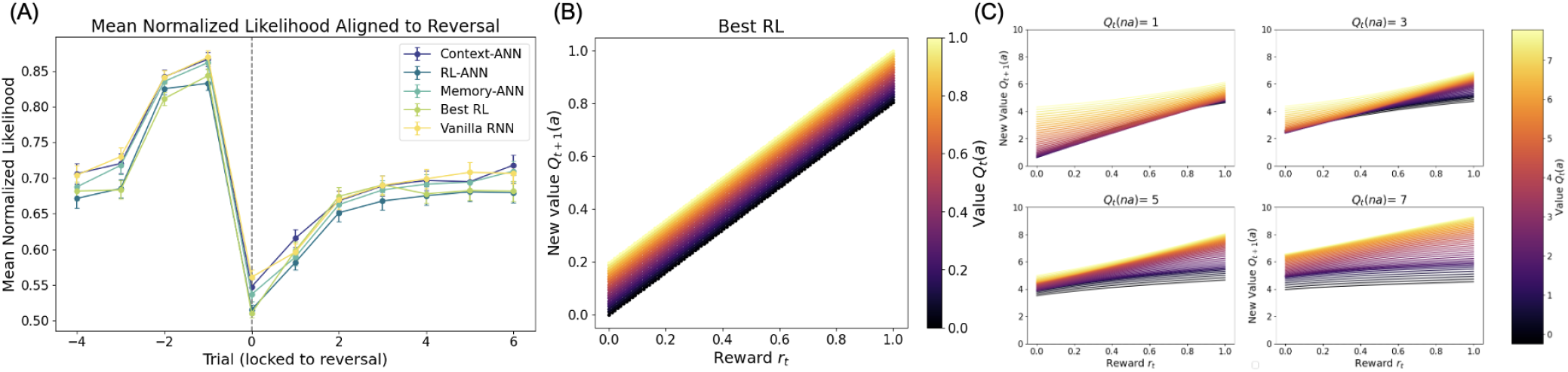
Value updating functions of Best RL and Context-ANN in Eckstein data. (A) Mean normalized likelihood of each model on human held-out data, aligned to reversal trials. Context-ANN and Vanilla RNN show the largest predictive accuracy. (B) Linear updating function of classical RL model. (C) Nonlinear and context dependent value updating functions of Context-ANN.

## Discussion

The present study applied classical cognitive modeling, neural network modeling, and HybridRNN to behavioral data from a reversal learning task. We improved the predictive accuracy of RL models to a level comparable to ANNs through an interpretable hybrid modeling structure, though important behavioral differences—such as the post-reversal dip—were not fully captured. Given the advantage of combining neural networks with cognitive modeling, the improvement in model prediction relied less on handcrafting and more on an automatic approach. Most importantly, HybridRNN enabled direct inspection of the trajectory of value drift, providing a platform to probe the value updating strategy that supports flexible learning. Our results suggest that Context-ANN provides the closest approximation to human behavior among the models considered. Further model inspection revealed nonlinear, context-dependent value updating functions, as well as structured hidden dynamics that reorganize around reversals, offering a promising approximation of the strategies supporting flexible adaptation.

The Context-ANN model in the present study suggested that context information is critical to human flexible reversal learning, differing from previous studies that applied HybridRNN to four-arm bandit tasks, where memory information was identified as the most important factor (Eckstein et al., 2022). Although adding memory input to RL-ANN improved its predictive accuracy for reversal learning behavior, its effect was not as good as that of context information. One possible reason is the different number of bandits used in this task, as Collins & Frank (2018) suggested that working memory plays a crucial role as the number of bandits increases. The two-armed bandit reversal learning task in the present study likely required a lower level of working memory to track bandit values. Furthermore, due to unpredictable switches in reward probabilities, the human brain recruits the prefrontal cortex to adapt to the changing environment rather than relying solely on the hippocampus (Guise & Shapiro, 2017), suggesting a potential difference in behavioral strategies. Previous pharmacological studies have also shown that dopamine manipulation improved short-term memory but impaired reversal learning (Mehta et al., 2001). Thus, over-reliance on memory in tasks with fewer choice options could negatively impact flexible learning behavior. Additionally, recent studies have interpreted reversal learning as a form of hidden state inference (e.g., Zika et al., 2023). From this perspective, the advantage of Context-ANN may stem from its capacity to approximate such inference mechanisms, as it integrates information from both choice options to infer the current latent state (Mishchanchuk et al., 2024). While not explicitly modeling belief states, this implicit approximation may bring its behavior closer to the strategy humans use to solve the task. This aligns with the Hierarchical Gaussian Filter (Mathys et al., 2011), which treat reversal learning as inference over latent states. Future comparisons with such models could help clarify the computational basis of flexible behavior. These might be possible reasons why Context-ANN outperformed Memory-ANN in predictive accuracy, on the current task, but not in the previous one with more choice options.

The flexibility of Context-ANN enabled us to examine how value updating strategies vary across different reward magnitudes and choice values. Previous studies have highlighted the critical role of contextual information in value updating (Palminteri et al., 2015) and the presence of nonlinearity in value updating functions (Nassar et al., 2019; Ji-An et al., 2023). However, a hybrid modeling approach—which integrates classical cognitive modeling with neural network modeling—can effectively reveal and quantify how nonlinear value updating strategies are modulated by contextual factors. Specifically, Context-ANN demonstrated that as the value of the unchosen action increases, the chosen action value becomes less sensitive to reward. This changing relationship with reward may be linked to participants’ attention allocation, as attention during reversal learning has been shown to be directed toward higher-value options (Oemisch et al., 2017). A higher unchosen action value could potentially reduce the attention allocated to the chosen action, thereby diminishing the extent to which its value is updated based on received rewards.

To better interpret Context-ANN, we analyzed the hidden dynamics within Context-ANN model (Mante et al., 2013; Flesch et al., 2022). Our results revealed structured activity patterns aligned with task-relevant variables, such as value differences and reversal points. These findings suggest that even in recurrent models with relatively simple architectures, internal representations can organize flexibly in response to changing task demands. This supports the idea that hybrid models can offer both predictive power and interpretable internal mechanisms, providing a valuable bridge between cognitive theory and neural network models.

A key limitation of the present study is its relatively small sample size (*n*_*trial*_ = 53, 520). Given the large number of parameters in ANN-based models, a larger dataset is generally required for reliable estimation (Alwosheel et al., 2018; Eckstein et al., 2024). The current sample size may not provide sufficient power to detect a statistically significant difference between the normalized likelihood of cognitive and neural network Models. Consequently, we relied on AIC scores as the primary model comparison metric, despite normalized likelihood is a more robust measure for cross-validation (Ji-An et al., 2023). In addition, the limited complexity of the task—specifically its two-option structure and fully anticorrelated reward schedule—may also reduce behavioral variability, making it more difficult to distinguish between competing models. Future work should consider larger and more behaviorally rich datasets—e.g., involving more choice options or stochastic reward structures—to better evaluate and distinguish competing computational accounts of learning behavior.

## Supplementary Analysis and Results

### Details of Hidden Unit Dynamics Analysis

To assess whether the model’s internal states encode contextual value information, we trained a logistic regression classifier to decode whether *Q*_A_ *> Q*_B_ on each trial. We used trials around reversal trials, and evaluated decoding accuracy using 5-fold cross-validation.

Two types of input features were tested: (1) the full hidden unit activations (8 dimensions), and (2) the top three principal components (PCs) of the hidden states. All features were z-scored. The classifier was implemented using `scikit-learn`‘s logistic regression with 1000 iterations. Decoding accuracy was 0.788 using the hidden units and 0.764 using the top 3 PCs.

Next, to examine whether the model’s internal dynamics exhibit attractor-like clustering, we applied dimensionality reduction and clustering to the hidden unit dynamics. The resulting PCA projections were standardized and clustered using the DBSCAN algorithm (`eps` = 0.5, `min_samples` = 10). This analysis revealed 4 distinct clusters in the low-dimensional space (see Fig. 6A), along with a small number of unassigned noise points. These attractor-like clusters suggest that the network organizes its internal dynamics into discrete regimes, potentially reflecting different value contexts during learning.

To quantify changes in internal state dynamics around reversals, we examined transitions between DBSCAN-identified clusters in the PCA-projected hidden state space. We tracked cluster membership transitions across adjacent trials, aligned to the reversal index (switch ∈ [− 7, +10]). For each transition (e.g., − 2→− 1), we computed the number of total transitions and the proportion that involved a change in cluster identity. This allowed us to estimate the likelihood of switching attractor states around reversal points. A detailed breakdown of cluster-to-cluster transitions is provided in Table5, and overall transition dynamics are shown in Figure 6B.

Overall, the proportion of inter-cluster transitions markedly increased following the reversal. While the transition rate remained relatively stable before reversal (mean change rate ≈ 36%), it sharply increased to over 40% during the first few trials post-reversal (e.g., 0→1: 42.06%, 1→2: 40.83%). This suggests that the Context-ANN reorganized its internal dynamics to adapt to the new environment, consistent with the notion of attractor reconfiguration.

Importantly, different types of transitions peaked at different time points. Transitions such as C0→C1 and C2→C3 showed an immediate increase at the first trial after reversal (trial 0), whereas C3→C0 and C1→C2 peaked later (around trial 1–2). This temporal asymmetry indicates that different internal reconfigurations unfold over distinct timescales, possibly reflecting a two-stage adaptation process: an initial detection and destabilization of outdated representations, followed by the formation of new task-aligned attractor states.

Taken together, these findings highlight the structured and temporally dynamic nature of internal state transitions in Context-ANN, supporting its ability to flexibly reorganize representations in response to environmental changes.

### Dataset from Eckstein et al., (2022)

We analyzed behavioral data from a probabilistic reversal learning task published by Eckstein et al. (2022). In this task, participants chose between two boxes on each trial in order to collect golden coins (i.e., rewards), which were probabilistically associated with the two options. The correct box was rewarded on 75% of trials, while the incorrect box never delivered rewards. Task contingencies switched unpredictably and without explicit cues, encouraging participants to track changes in the underlying reward structure. Reversals were only allowed after participants had collected a random number of rewards (between 7 and 15), and the first correct choice after a switch was always rewarded. Participants completed 120 trials, preceded by a tutorial phase that familiarized them with task rules and reversals.

For the present study, we followed the original exclusion criteria to remove participants with low task engagement or missing demographic information. After exclusion, the final sample comprised 179 adolescents (under age 18; 96 males and 83 females), 57 undergraduates aged 18–28 (19 males and 38 females), and 55 adult community participants (26 males and 29 females), yielding a total sample size of 291 participants for model fitting and analysis.

### Quantitative and Qualitative Model Comparison on Eckstein Dataset

To evaluate the robustness of our findings, we trained and tested the same series of cognitive models, RNNs, and HybridRNN models on the independent dataset from Eckstein et al. (2022). As shown in Table 6 and Table 7, we successfully replicated the main pattern observed in our primary dataset: replacing the hand-crafted value update function in traditional cognitive models with a neural network significantly improved model fit to human behavior. Notably, Context-ANN once again achieved predictive accuracy on par with the Vanilla RNN, further demonstrating that hybrid models can combine strong predictive performance with improved interpretability.

In addition, following a reviewer’s suggestion, we performed a trial-wise normalized likelihood analysis aligned to the reversal point (see Fig. 7A). This analysis provides a more intuitive view of model predictions across learning transitions. We found that both Context-ANN and Vanilla RNN consistently achieved the highest likelihood across trials compared to other models, indicating strong predictive alignment with human behavior. However, all models exhibited a notable drop in prediction likelihood at the exact reversal trial, highlighting a limitation in capturing human switching dynamics.

One possible reason is that the Eckstein dataset spans a wide developmental age range, including children, adolescents, and adults. As previous research suggests, adolescence is a period of rapid and heterogeneous changes in learning and decision-making strategies. This heterogeneity may pose challenges for models like HybridRNNs that rely on shared trial-level patterns across participants. Therefore, we caution against overinterpreting this dataset as a benchmark for model comparison. Future work may benefit from identifying or generating datasets with more homogeneous populations and better-controlled task environments for model discovery and validation.

### Information Processing

We replicated the context-dependent value updating pattern in the Eckstein dataset (see Fig.7C). A linear regression predicting *Q*_*t*+1_(*a*) revealed a significant negative interaction between *r*_*t*_ and *Q*_*t*_(*na*) (β =−0.28, *p <* .001; Table 8), confirming that rewards were weighted more strongly when the unchosen option had lower expected value.

**Table 6:**
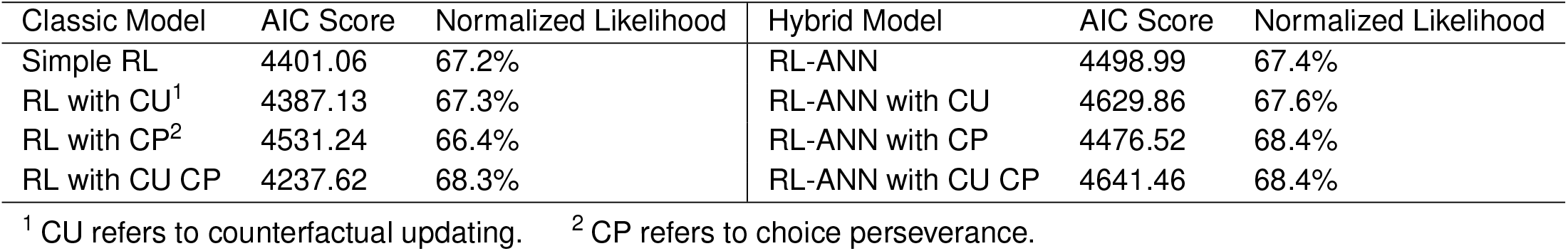
Quantitative model comparisons of cognitive and hybrid model series on Eckstein dataset.

| Classic Model | AIC Score | Normalized Likelihood | Hybrid Model | AIC Score | Normalized Likelihood |
| --- | --- | --- | --- | --- | --- |
| Simple RL | 4401.06 | 67.2% | RL-ANN | 4498.99 | 67.4% |
| RL with CU<sup>1</sup> | 4387.13 | 67.3% | RL-ANN with CU | 4629.86 | 67.6% |
| RL with CP<sup>2</sup> | 4531.24 | 66.4% | RL-ANN with CP | 4476.52 | 68.4% |
| RL with CU CP | 4237.62 | 68.3% | RL-ANN with CU CP | 4641.46 | 68.4% |

<sup>1</sup> CU refers to counterfactual updating. <sup>2</sup> CP refers to choice perseverance.

**Table 7:**
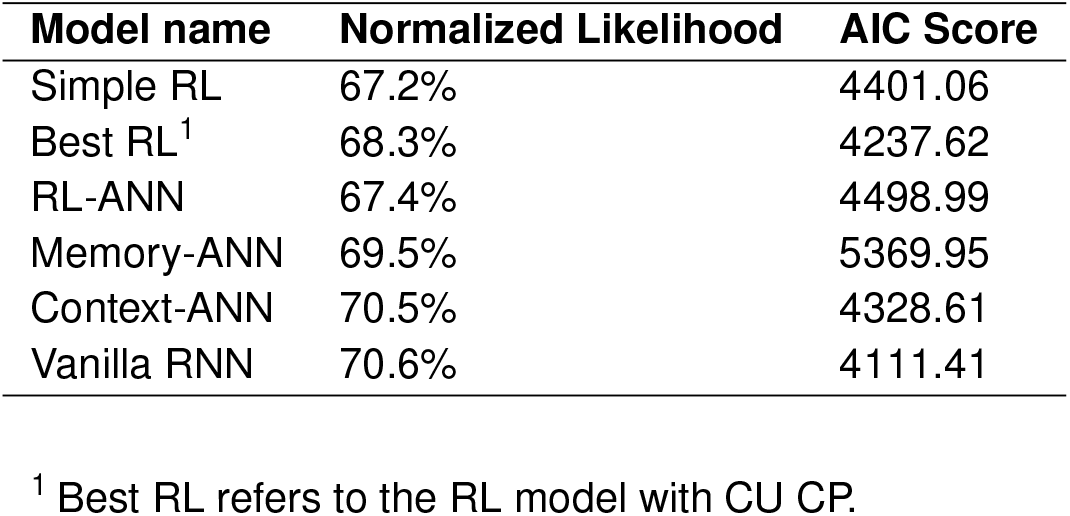
Quantitative model comparisons of cognitive models, RNN, and HybridRNN models on Eckstein dataset.

| Model name | Normalized Likelihood | AIC Score |
| --- | --- | --- |
| Simple RL | 67.2% | 4401.06 |
| Best RL<sup>1</sup> | 68.3% | 4237.62 |
| RL-ANN | 67.4% | 4498.99 |
| Memory-ANN | 69.5% | 5369.95 |
| Context-ANN | 70.5% | 4328.61 |
| Vanilla RNN | 70.6% | 4111.41 |

<sup>1</sup> Best RL refers to the RL model with CU CP.

**Table 8:**
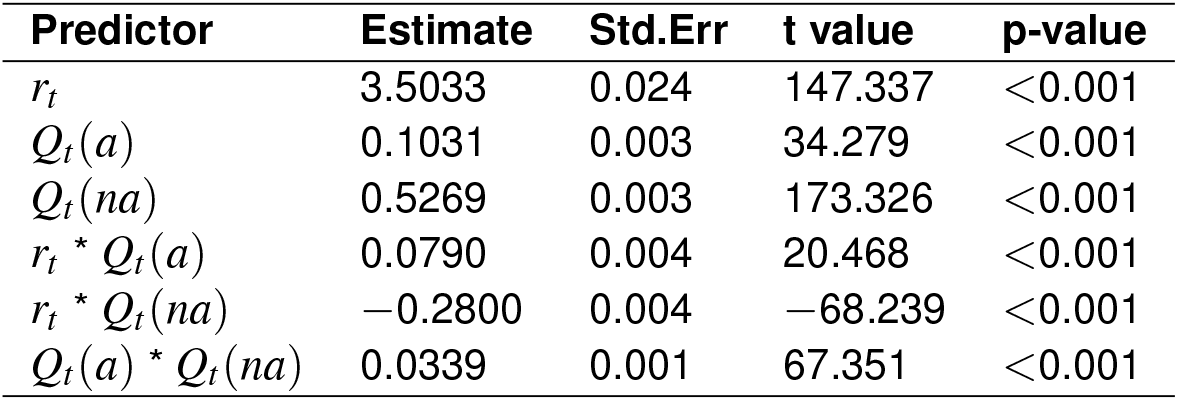
Linear regression results on predicting value of chosen action on Eckstein dataset.

| Predictor | Estimate | Std.Err | t value | p-value |
| --- | --- | --- | --- | --- |
| $r_t$ | 3.5033 | 0.024 | 147.337 | <0.001 |
| $Q_t(a)$ | 0.1031 | 0.003 | 34.279 | <0.001 |
| $Q_t(na)$ | 0.5269 | 0.003 | 173.326 | <0.001 |
| $r_t * Q_t(a)$ | 0.0790 | 0.004 | 20.468 | <0.001 |
| $r_t * Q_t(na)$ | -0.2800 | 0.004 | -68.239 | <0.001 |
| $Q_t(a) * Q_t(na)$ | 0.0339 | 0.001 | 67.351 | <0.001 |

